# Thermal proteome profiling identifies mitochondrial aminotransferases involved in cysteine catabolism via persulfides in plants

**DOI:** 10.1101/2025.05.23.655777

**Authors:** Björn Heinemann, Jannis Moormann, Shivsam Bady, Cecile Angermann, Andrea Schrader, Tatjana M. Hildebrandt

## Abstract

Cysteine is a central metabolite in plant sulfur metabolism, with key roles in biosynthesis, redox regulation, and stress responses. While a mitochondrial cysteine degradation pathway has been described, the enzyme catalyzing its initial transamination step remained unidentified. Here, we applied thermal proteome profiling (TPP) to Arabidopsis mitochondria to uncover cysteine-interacting proteins. TPP successfully detected known cysteine-utilizing enzymes, validating its utility in plant metabolic research. Among newly identified targets were two aminotransferases annotated as alanine and aspartate aminotransferases that catalyze the transamination of cysteine to 3-mercaptopyruvate *in vitro*. These enzymes, together with the sulfurtransferase STR1 and the persulfide dioxygenase ETHE1, reconstituted a complete mitochondrial cysteine catabolic pathway. Kinetic data indicate that alanine aminotransferase, in particular, may function *in vivo* under physiological cysteine levels. Additionally, GABA aminotransferase was inhibited by cysteine, suggesting a regulatory role in stress metabolism. Beyond enzyme identification, the dataset provides a resource for exploring cysteine-mediated regulation of transporters, RNA-editing factors, and respiratory components. Given cysteine’s emerging role as a metabolic signal in stress responses, and the importance of allosteric regulation in amino acid metabolism, these findings highlight the broader regulatory potential of cysteine–protein interactions in plants. This study demonstrates the utility of TPP for elucidating metabolite-protein networks and advancing our understanding of plant mitochondrial metabolism.

## Introduction

The amino acid cysteine holds a central position in plant sulfur metabolism, serving as the entry point of reduced sulfur into organic molecules. It is a precursor for a wide range of sulfur-containing compounds including methionine, cofactors such as thiamin, lipoic acid, biotin, Fe-S clusters, and the molybdenum cofactor (Droux, 2004; Giovanelli et al., 1985; van Hoewyk et al., 2008). The tripeptide glutathione (γ-glutamyl-cysteinyl-glycine) represents a major antioxidant and redox buffer in plants, protecting cells from oxidative damage and regulating cellular signaling pathways. Its antioxidant function is rooted in the redox potential of the thiol group (Foyer & Noctor, 2011). In addition to its role in oxidative stress protection, glutathione contributes to heavy metal and xenobiotic detoxification, as well as to defense responses against pathogens (Noctor et al., 2024). Phytochelatins, synthesized from glutathione-derived cysteine, play a crucial role in heavy metal detoxification by chelating toxic metals and facilitating their sequestration in vacuoles (Cobbett & Goldsbrough, 2002). Cysteine also serves as a precursor for the biosynthesis of various secondary metabolites, such as glucosinolates and camalexin, which play important roles in plant defense against herbivores and pathogens (Halkier & Gershenzon, 2006; Su et al., 2011). Cysteine residues in proteins support structure, catalysis, and redox regulation through disulfide bond formation as well as reversible thiol modifications such as glutathionylation, nitrosylation, and persulfidation (Begara-Morales et al., 2016; Buchanan & Balmer, 2005; Moseler et al., 2024).

Cysteine is synthesized via the sulfur assimilation pathway. Sulfate taken up from the soil is activated by ATP sulfurylase and reduced via sulfite to sulfide by APS reductase and sulfite reductase, respectively. O-acetylserine (OAS), synthesized from serine and acetyl-CoA by serine acetyltransferase (SERAT), serves as the carbon skeleton for cysteine biosynthesis. The final step, incorporation of sulfide into OAS to form cysteine, is catalyzed by OAS-(thiol)lyase (OASTL) (Takahashi, 2010; Takahashi et al., 2011). SERAT and OASTL form a regulatory cysteine synthase complex that adjusts synthesis in response to substrate availability (Droux, 2003; Wirtz & Hell, 2006). Isoforms of both enzymes are present in the cytosol (SERAT1;1, SERAT3;1, SERAT3;2; OASTL-A), plastids (SERAT2;1; OASTL-B), and mitochondria (SERAT2;2; OASTL-C), enabling localized cysteine synthesis across compartments (Hell & Wirtz, 2011; Ruffet et al., 1995; Watanabe et al., 2008). The cytosol and plastids account for 95% of total OASTL activity and null mutant analyses suggest a certain degree of functional redundancy among the major isoforms (Birke et al., 2013; Heeg et al., 2008; Krüger et al., 2009). However, evidence is accumulating for the functional relevance of compartment-specific cysteine metabolism.

Mitochondrial cysteine synthesis provides substrates for the production of cysteinyl-tRNA during translation of mitochondrial-encoded proteins but also supports other compartment-specific functions (Fig. 1). Cysteine serves as a sulfur donor for the formation of Fe-S clusters, which are critical cofactors for several mitochondrial enzymes, required for the biosynthesis of lipoic acid and biotin and can also be exported to the cytosol (Couturier et al., 2013). The capacity for cysteine synthesis in the mitochondrial matrix is also necessary for local detoxification of the potent cytochrome c oxidase inhibitors cyanide and hydrogen sulfide. Cyanide, a byproduct of ethylene and camalexin biosynthesis in Arabidopsis, is detoxified by cyanoalanine synthase (CAS-C1), which uses cysteine as a substrate to convert cyanide into β-cyanoalanine and hydrogen sulfide (Böttcher et al., 2009; Hatzfeld et al., 2000; Peiser et al., 1984; Watanabe et al., 2008). Accumulating hydrogen sulfide is then re-assimilated into cysteine by OASTL-C, completing a local detoxification cycle. Thus, mitochondrial cysteine metabolism is essential for maintaining electron transport chain activity (Cooper & Brown, 2008). Beyond detoxification and biosynthesis, cysteine participates in regulatory processes with compartment specific mechanisms and functions. Recent work has demonstrated that cysteine triggers defense responses in Arabidopsis and enhances resistance to *Pseudomonas syringae*, suggesting a signaling function beyond metabolic necessity. The balance between cytosolic and organellar cysteine synthesis appears critical for proper immune function (Moormann et al., 2025). Post-translational modifications, such as disulfide bond formation and glutathionylation, nitrosylation, and persulfidation, mediate signaling processes by regulating protein function and activity (Begara-Morales et al., 2016; Buchanan & Balmer, 2005; Moseler et al., 2024). The mechanism of protein persulfidation and de-persulfidation in the different subcellular compartments is largely unknown and the physiological relevance of this post-translational modification in plant mitochondria has not been established yet (Moseler et al., 2024). It might well be linked to the mitochondrial cysteine degradation pathway, which produces persulfides as intermediates (Fig. 1, Höfler et al., 2016). Cysteine catabolism in the mitochondria starts with a transamination step to 3-mercaptopyruvate, which then transfers its thiol group to glutathione catalyzed by mercaptopyruvate sulfurtransferase (STR1, AT1G79230) producing glutathione persulfide. Yeast mercaptopyruvate sulfurtransferase has recently been shown to act as a protein persulfidase, and it is tempting to speculate that this function might also be present and relevant in plant mitochondria (Pedre et al., 2023). To complete the cysteine catabolic pathway, glutathione persulfide can be oxidized to sulfite by the persulfide dioxygenase ETHE1 (AT1G53580) and converted to thiosulfate via addition of a second persulfide group by STR1 (Höfler et al., 2016; Krüßel et al., 2014). For more than 10 years after initial identification of the mitochondrial cysteine degradation pathway in plants the aminotransferase catalyzing the initial step has remained enigmatic.

**Fig. 1:**
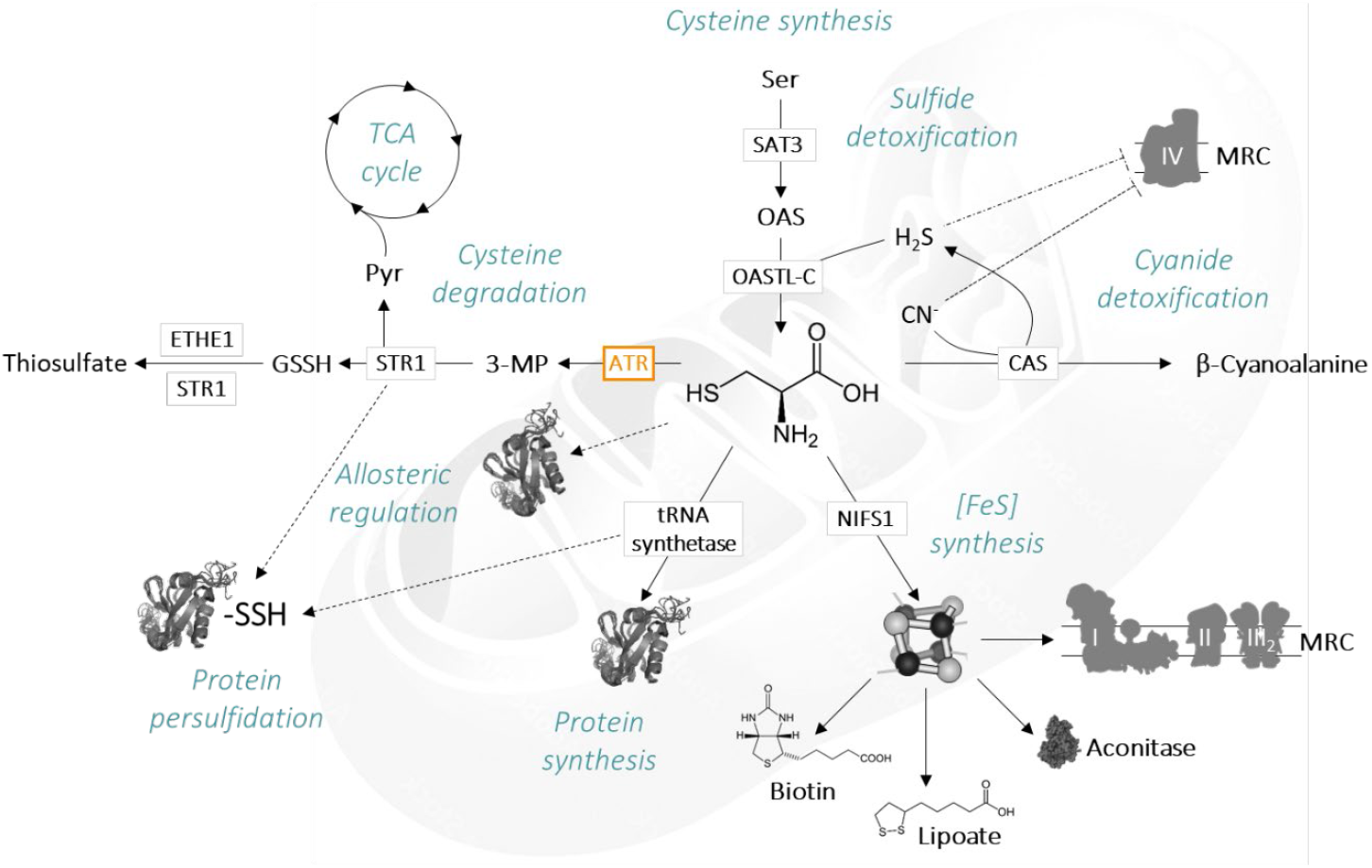
Mitochondrial cysteine metabolism. Plant mitochondria contain the enzymes required for cysteine synthesis and for the production of cysteinyl-tRNA for protein synthesis. In addition, cysteine serves as a sulfur donor during the synthesis of iron-sulfur clusters, which are required as cofactors for several mitochondrial enzymes and provide the sulfur for the cofactors biotin and lipoate. Cysteine is also involved in the detoxification of two different inhibitors of cytochrome c oxidase. It is a substrate of β-cyanoalanine synthase catalyzing cyanide detoxification, and cysteine synthesis detoxifies accumulating hydrogen sulfide. Regulatory functions can be mediated either by post-translational protein modification or allosteric effects. Mitochondrial cysteine catabolism proceeds via transamination to 3-mercaptopyruvate, subsequent transfer of the thiol group to glutathione by mercaptopyruvate sulfurtransferase, and oxidation to sulfite by the persulfide dioxygenase ETHE1 followed by transfer of an additional thiol group to produce thiosulfate. Mitochondrial aminotransferases catalyzing the first step of this cysteine degradation pathway have been identified in the frame of this study. ATR, aminotransferase; CAS, β-cyanoalanine synthase; ETHE1, persulfide dioxygenase; MRC, mitochondrial respiratory chain; NIFS1, cysteine desulfurase; OASTL-C, O-actylserine(thiol)lyase C; SAT3, serine acetyltransferase 3; STR1, mercaptopyruvate sulfurtransferase; TCA cycle, tricarboxylic acid cycle.

We here explore the potential of thermal proteome profiling (TPP) as a new powerful tool to screen for mitochondrial cysteine metabolic enzymes. TPP is based on the principle that ligand binding affects the thermostability of proteins. Depending on the nature of the interaction proteins can be stabilized but also destabilized during interaction with small molecules. TPP identifies these changes in the melting temperatures of proteins that are able to bind a metabolite of interest. Aliquots of a protein extract are heated to different temperatures in the presence and absence of potential ligands. Afterwards, precipitated proteins are removed by centrifugation, and the relative abundance of each protein in the remaining non-denatured fraction is determined by mass spectrometry (Fig. 2). While originally developed for drug target discovery in biomedical research (Savitski et al., 2014), the application of TPP in plant systems remains largely unexplored. Here, we demonstrate its effectiveness in uncovering a metabolic enzyme involved in plant primary metabolism, a mitochondrial cysteine aminotransferase. We identify two mitochondrial aminotransferases, previously annotated as aspartate and alanine aminotransferases, that use cysteine as an amino donor and 3-mercaptopyruvate as a keto acid acceptor *in vitro*, placing them as functional components of the mitochondrial cysteine oxidation pathway. TPP thus proved successful in filling a critical gap in our understanding of mitochondrial sulfur catabolism and its integration into cellular metabolism and signaling. Beyond this, the dataset reveals a broader set of cysteine-binding proteins, providing a resource for further exploration of cysteine metabolism and its regulatory roles in plant mitochondria.

**Fig. 2:**
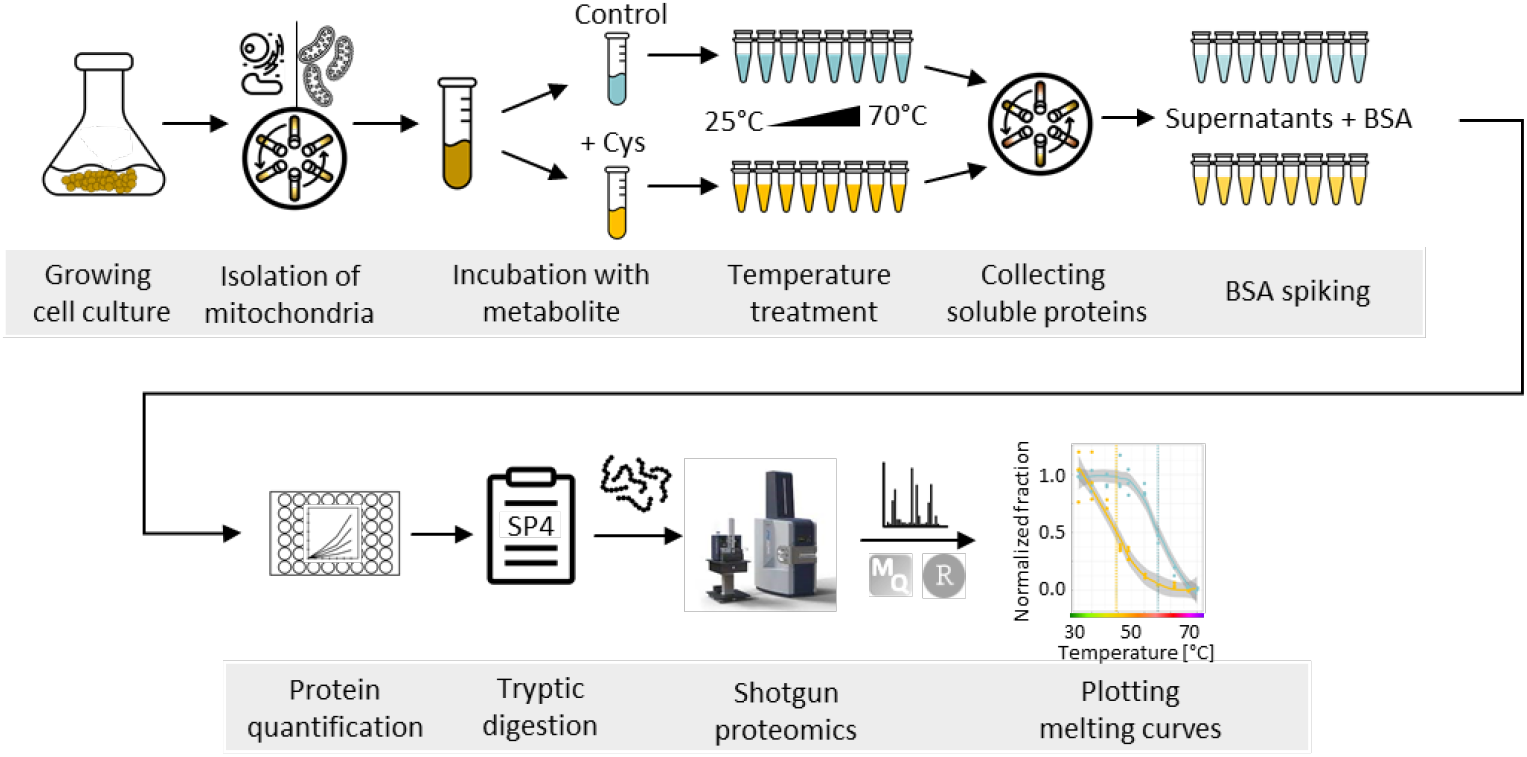
Thermal proteome profiling (TPP) workflow. Mitochondria were isolated from *Arabidopsis thaliana* cell culture via differential and density gradient centrifugation. The mitochondrial extract was split and either incubated with L-cysteine or water. The treated extracts were then aliquoted for the temperature treatment (n=3). Treated aliquots were centrifuged and supernatants were collected. BSA was added as internal standard to preserve the information of differences in protein concentrations. Protein concentrations were then adjusted to 1 µg µl^-1^ to ensure equal conditions during tryptic digestion. The resultant peptides were quantified using shotgun mass spectrometry, MaxQuant software and a custom R-script.

## Results

### Thermal proteome profiling of the *Arabidopsis thaliana* mitochondrial proteome

Thermal mitochondrial-proteome profiling requires protein samples with sufficient coverage and concentration of the target proteome to allow reliable melting curve generation. To this end, mitochondria were isolated from *Arabidopsis thaliana* cell suspension cultures, yielding a protein fraction highly enriched in mitochondrial proteins (88%). Aliquots of this fraction were subjected to a temperature gradient from 25°C to 70°C in the presence or absence of 1 mM L-cysteine, each in triplicate. Soluble proteins were identified and quantified via shotgun proteomics (Fig. 2), resulting in the detection of 3903 protein groups.

Principal component analysis (PCA) clearly separated the samples along the temperature gradient, reflecting progressive thermal denaturation of individual proteins (Fig. 3A). Correspondingly, the average normalized abundance of mitochondrial proteins decreased gradually with increasing temperature (Fig. 3B). Melting curves were generated from normalized abundance data using a custom R script (see Methods). We applied stringent filters to exclude proteins detected in fewer than two replicates at 25°C, non-mitochondrial proteins, and proteins with inconsistent melting curves due to high variability within the dataset (see Suppl. Fig. S1 for filtering workflow).

**Fig. 3:**
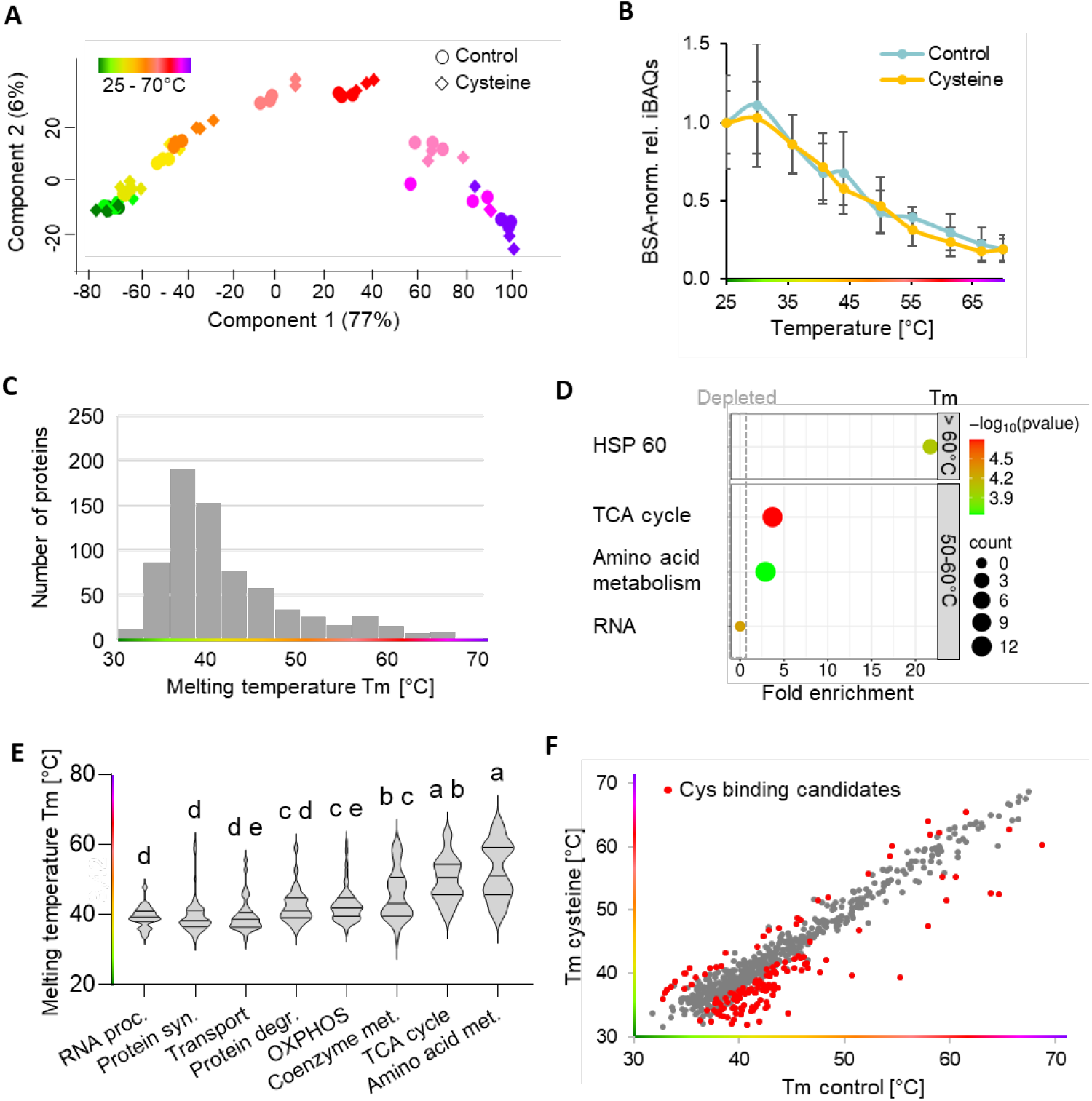
Thermal proteome profiling of the *Arabidopsis thaliana* mitochondrial proteome. **A** iBAQ-based principal component analysis of the proteomics dataset. **B** Mean normalized soluble fractions of all quantified mitochondrial proteins (BSA-normalized and relative to 25°C). **C** Histogram illustrating the distribution of melting temperatures in the mitochondrial proteome. **D** Enrichment analysis of highly and extremely heat stable mitochondrial protein fractions. **E** Mean thermostability of proteins involved in major mitochondrial pathways. **F** Melting temperatures (Tm) of the 716 mitochondrial proteins with high-quality melting curves. Candidates for cysteine-interacting proteins (= 146 proteins showing melting point shifts in the presence of 5 mM cysteine) are highlighted in red.

After filtering, high-confidence melting curves were obtained for 716 mitochondrial proteins (Suppl. Dataset S1). Calculated melting temperatures (Tm) ranged from 31.7°C to 68.7°C, with most proteins denaturing around 40°C (Fig. 3C). Notably, a subset of proteins displayed exceptional thermal stability: 12.2% had Tm values between 50°C and 60°C, and 4.2% above 60°C. The thermotolerant fraction (Tm = 50–60°C) was significantly enriched in enzymes of the tricarboxylic acid (TCA) cycle and amino acid metabolism (Fig. 3D), while the extremely heat-stable proteins (Tm > 60°C) included additional enzymes of amino acid metabolism and a group of heat shock proteins. Mean thermostability also varied significantly across the major mitochondrial pathways (Fig. 3E). Proteins involved in amino acid metabolism and the TCA cycle were generally more heat-stable, whereas RNA editing factors exhibited low melting temperatures and were absent from the thermotolerant fractions.

### Cysteine-induced thermal stability shifts in the *Arabidopsis thaliana* mitochondrial proteome

In this study, we applied TPP to identify mitochondrial proteins that interact with L-cysteine. Comparison of melting curves between cysteine-treated and control samples revealed significant thermal stability shifts for 146 mitochondrial proteins (Fig. 3F). Among these were three enzymes known to utilize L-cysteine as a substrate (Fig. 1), the cysteine desulfurase NifS1 (AT5G65720, ΔTm = -2.9°C), O-actylserine-(thiol)lyase C (OASTL-C; AT3G59760; ΔTm = -4.1°C), and cysteinyl-tRNA synthetase (AT2G31170, ΔTm = 6.1°C) (Suppl. Dataset S1). These findings serve as a positive control, validating the suitability of TPP for identifying metabolic enzymes that use a specific compound of interest as a substrate. In addition, our dataset offers a resource for identifying potential targets of cysteine-mediated metabolic regulation including transporters, RNA editing factors, and respiratory chain components (Suppl. Dataset S1).

### Identification and biochemical characterization of mitochondrial cysteine aminotransferase candidates

Among the cysteine-interacting mitochondrial proteins five were annotated as aminotransferases (Fig. 4, Suppl. Dataset S1). Depending on the site and nature of interaction this could either be candidates for a mitochondrial cysteine aminotransferase catalyzing transamination of cysteine to 3-mercaptopyruvate or enzymes regulated by cysteine. The candidate aminotransferases were annotated as gamma-aminobutyric acid (GABA) aminotransferase (GABA-AT, AT3G22200, ΔTm = -10.9°C), aspartate aminotransferase (Asp-AT, AT2G30970, ΔTm = -8.2°C), alanine-glyoxylate aminotransferase (AT4G39660, ΔTm = -11.2°C), and alanine aminotransferases (Ala-AT, AT1G17290, ΔTm = 3.2°C; AT1G72330, ΔTm = 3.5°C). The first three candidates were strongly destabilized in the presence of cysteine whereas the alanine aminotransferases were stabilized (Fig. 4).

**Fig. 4:**
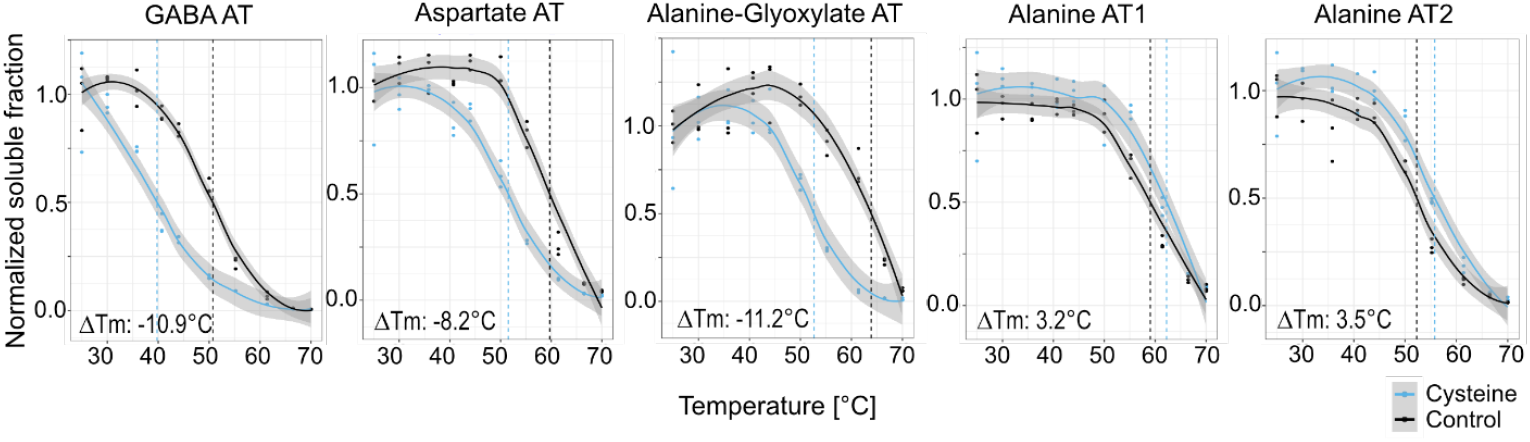
Candidates for a mitochondrial cysteine aminotransferase identified by TPP. Melting curves of the five mitochondrial aminotransferases (AT) showing a shift in the melting point (marked by dotted lines) in the presence of 5 mM cysteine. Grey shadows: Locally estimated scatterplot smoothing (LOESS); GABA aminotransferase (AT3G22200), Aspartate aminotransferase 1 (AT2G30970), Alanine-glyoxylate aminotransferase 2 (AT4G39660), Alanine aminotransferase 1 (AT1G17290), Alanine aminotransferase 2 (AT1G72330).

For functional analysis and biochemical characterization, the genes (excluding predicted mitochondrial targeting sequences) were expressed in *Escherichia coli*, and the His-tagged proteins were purified via Ni-NTA affinity and size-exclusion chromatography (Suppl. Fig. S2, S3). Despite repeated attempts to optimize the protocol, active alanine-glyoxylate aminotransferase could not be obtained, consistent with previous reports (Liepman & Olsen, 2003). In contrast, GABA-AT, Asp-AT, and Ala-AT were successfully expressed and purified, with Ala-AT1 (AT1G17290) selected as the representative isoform for further analysis.

Enzyme assays confirmed that the purified aminotransferases were catalytically active with their canonical substrates and the kinetic parameters were in the expected range (Fig. 5, Table 1, Suppl. Fig. S4). Ala-AT and Asp-AT also accepted L-cysteine as an amino acid substrate, albeit with reduced catalytic efficiency. (Fig. 5A, B; Table 1). However, kinetic analysis of Ala-AT revealed a K_m_-value of 0.77 ± 0.10 mM for L-Cys compared to 2.51 ± 0.18 mM for L-Ala indicating a higher affinity of the enzyme for cysteine as an amino group donor (Fig. 5A, Table 1). In contrast, Asp-AT showed a higher affinity for its established substrate with K_m_-values of 3.33 ± 0.06 mM for L-Asp vs. 15.14 ± 0.64 mM for L-Cys (Fig. 5B, Table 1). GABA-AT did not metabolize cysteine in our experimental setup but was inhibited by it. Kinetic evaluation suggested a competitive mode of inhibition with a K_i_ of 0.42 mM Cys for transamination of GABA and 0.18 mM Cys for the reverse reaction using Ala as an amino group donor (Fig. 5C).

**Table 1:**
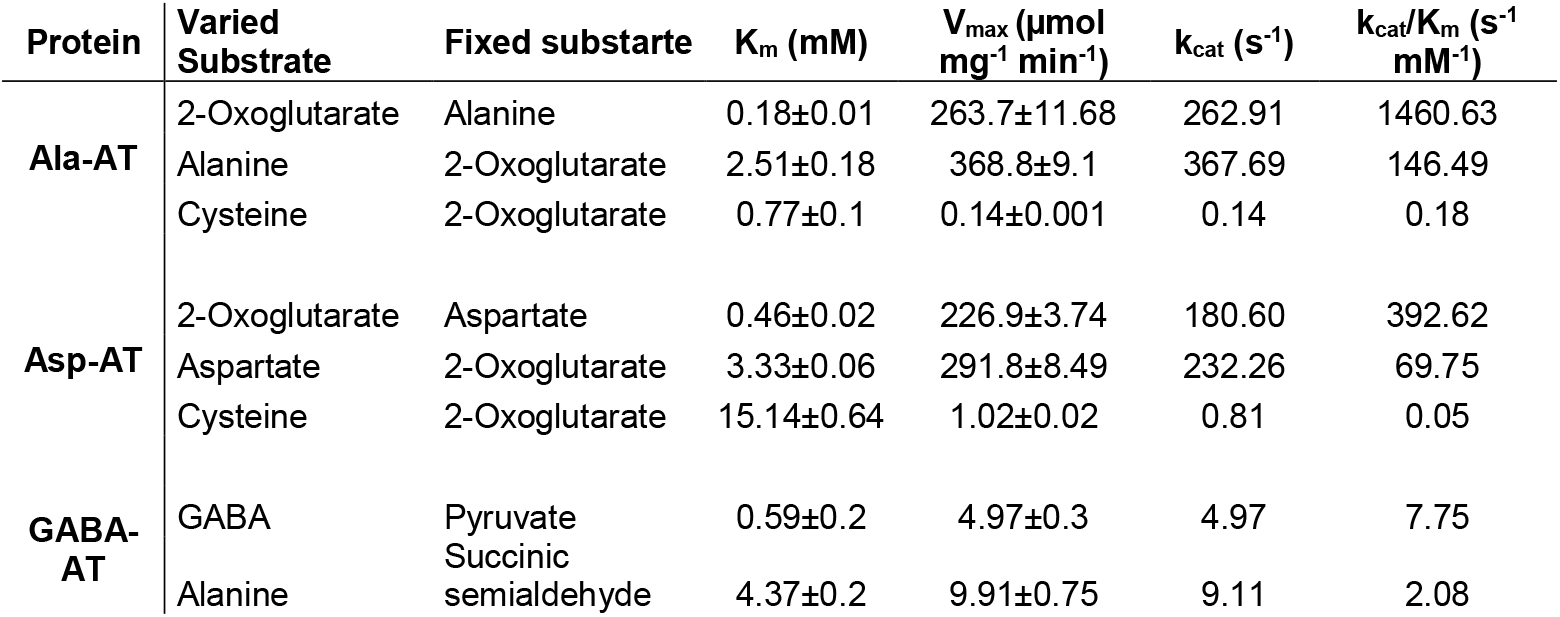
Kinetic parameters for the purified recombinant Ala-AT, Asp-AT and GABA-AT.

**Fig. 5:**
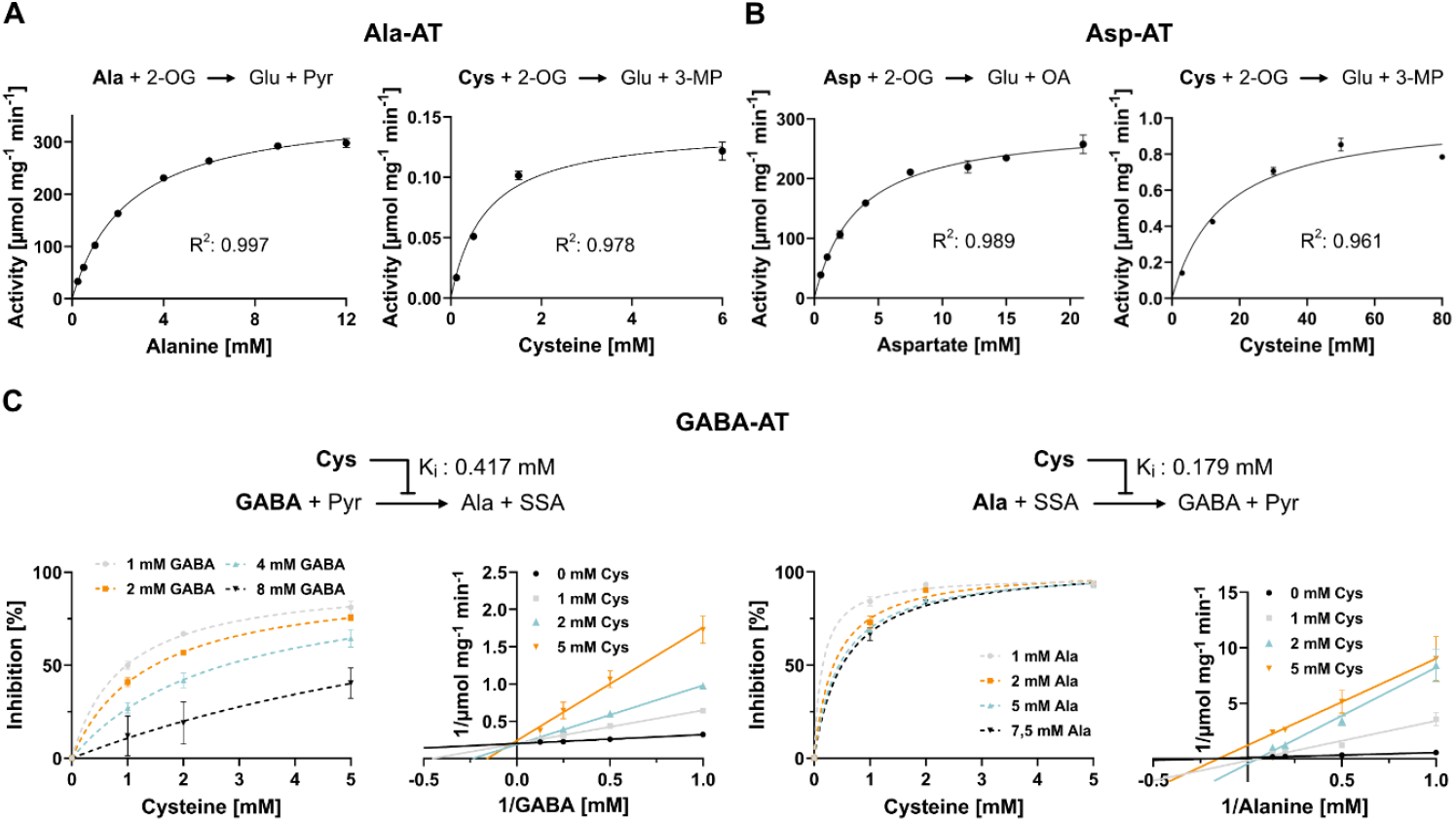
Biochemical characterization of mitochondrial cysteine aminotransferase candidates. **A** Michaelis Menten kinetics for alanine aminotransferase 1 (AT1G17290) with L-Ala and L-Cys as a variable substrate. **B** Michaelis Menten kinetics for aspartate aminotransferase 1 (AT2G30970) with L-Asp and L-Cys as a variable substrate. **C** Inhibition kinetics for GABA aminotransferase 1 (AT3G22200) by L-Cys for both directions of the reaction. 2-OG, 2-oxoglutarate; 3-MP, 3-mercaptopyruvate; GABA, gamma-aminobutyric acid; OA, oxaloacetate; Pyr, pyruvate; SSA, succinic semialdehyde.

### Reconstruction of the complete plant mitochondrial cysteine catabolic pathway

We next tested, whether the candidate aminotransferases could support the proposed mitochondrial cysteine catabolic pathway. The pathway involves four enzymatic steps catalyzed by three enzymes: transamination of L-cysteine to 3-mercaptopyruvate, conversion to glutathione persulfide by STR1 (AT1G79230), oxidation to sulfite by the persulfide dioxygenase ETHE1 (AT1G53580), and transfer of an additional persulfide to sulfite by STR1 producing the final product thiosulfate (Fig. 6A). All components were heterologously expressed in and purified from *E. coli*. (Suppl. Fig. S5). ETHE1 activity was measured by oxygen consumption and confirmed via HPLC-based sulfite quantification (Fig. 6A, 6F). In the presence of STR1, ETHE1-dependent oxygen consumption could be triggered using 3-mercaptopyruvate as substrate, leading to equimolar thiosulfate production (Fig. 6C, 6G). The complete three-enzyme system successfully catalyzed the oxidation of L-cysteine to thiosulfate when either Asp-AT or Ala-AT was included (Fig. 6D, 6E, 6H, 6I). Pathway activity nearly matched that of isolated ETHE1 when the aminotransferase and STR1 were provided in excess. Protein abundance estimates from the mitochondrial fraction revealed that Ala-AT, Asp-AT, and STR1 were present at about 10-fold higher levels than ETHE1, supporting the pathway’s feasibility *in vivo* (Suppl. Dataset S1). This is further supported by the higher spatial proximity of the enzymes in the mitochondrial matrix compared to the *in vitro* system. Consistent with its lack of cysteine-metabolizing activity, GABA-AT did not support ETHE1-mediated oxygen consumption from cysteine.

**Fig. 6:**
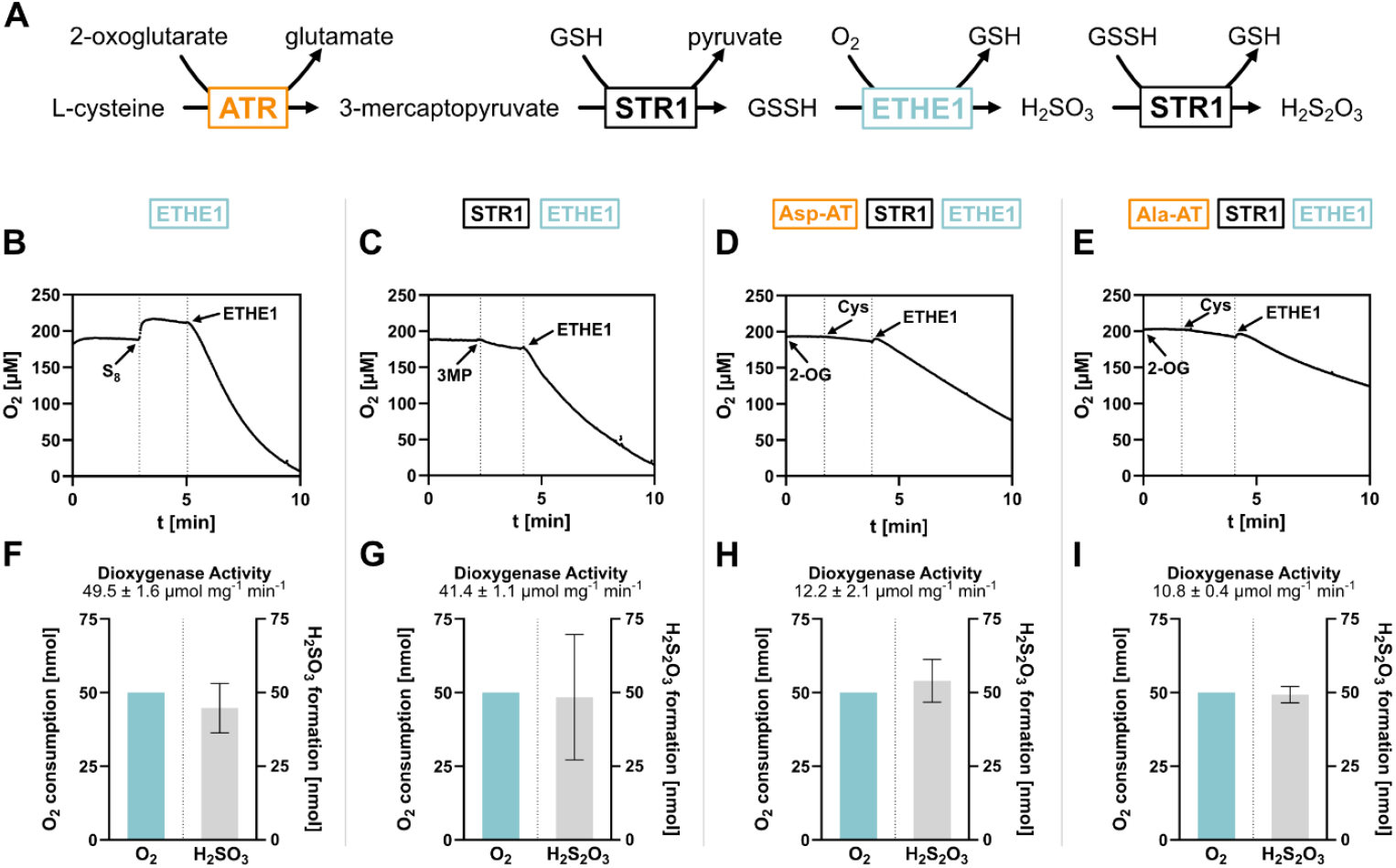
Reconstruction of the complete plant mitochondrial cysteine catabolic pathway *in vitro*. **A** Reaction scheme of the mitochondrial cysteine catabolic pathway. **B-E** Original traces of oxygen electrode measurements illustrating pathway activity based on oxygen consumption by the persulfide dioxygenase (ETHE1) step. The reactions contained GSH as well as the enzymes listed above, except for ETHE1, which was added at the indicated timepoint. Substrates were supplied as noted in the graph **F-I** Stoichiometry between oxygen consumption and product accumulation. Samples were taken from the reaction mix after exactly 50 nmol O_2_ had been consumed and thiol contents were analyzed by HPLC **B**,**F** Activity of the persulfide dioxygenase ETHE1 with GSSH (non-enzymatically produced from GSH plus elemental sulfur) as a substrate. **C**,**G** Activity of the partial cysteine catabolic pathway consisting of the sulfurtransferase STR1 plus ETHE1 with 3-mercaptopyruvate (3MP) as a substrate. **D**,**H** Activity of the complete cysteine catabolic pathway consisting of aspartate aminotransferase 1, STR1 and ETHE1 with cysteine as a substrate. **E**,**I** Activity of the complete cysteine catabolic pathway consisting of alanine aminotransferase 1, STR1 and ETHE1 with cysteine as a substrate.

To get a first impression about the metabolic integration of the three aminotransferases in mitochondrial sulfur metabolism as well as the physiological context of these networks we compared co-expression profiles available on ATTED (https://atted.jp/). There was a strong overlap of 43 genes between the top 100 co-expressed genes for STR1 and Asp-AT, and those were mainly involved in the mitochondrial housekeeping functions TCA cycle and oxidative phosphorylation (Suppl. Fig. S6A, Suppl. Dataset S2). The co-expression profile of ETHE1 was most similar to that of GABA-AT with 31 common genes in the top 100 (Suppl. Fig. S6B). In this case the group of overlapping genes was dominated by enzymes involved in lipid catabolism as well as amino acid transport and degradation, placing these enzymes (ETHE1 and GABA-AT) into the context of energy deficiency stress (Suppl. Dataset S2).

## Discussion

### Thermostability of plant mitochondrial proteins

Our thermal profiling of the *Arabidopsis thaliana* mitochondrial proteome revealed marked differences in the average thermostability of distinct functional categories. Notably, enzymes of the tricarboxylic acid (TCA) cycle and amino acid catabolism exhibited high melting temperatures, suggesting an intrinsic robustness that may be critical under conditions of energy limitation. During episodes of heat stress often coinciding with drought, plants frequently experience compromised photosynthesis and ATP depletion. Under such constraints, the persistence of a thermo-stable core of TCA cycle enzymes would help sustain mitochondrial ATP production when alternative energy-generating pathways falter. Similarly, the high thermal stability of amino acid catabolic enzymes likely reflects their central role in maintaining metabolic flexibility under stress. Amino acids act as alternative respiratory substrates, contributing intermediates to the TCA cycle and, in some cases, directly donating electrons to the mitochondrial electron transport chain via the ubiquinone pool (Hildebrandt et al., 2015). This catabolic route becomes especially critical during carbohydrate starvation, as demonstrated by the compromised stress tolerance and shortened survival of mutants deficient in key steps of amino acid degradation (Araújo et al., 2010; Hirota et al., 2018; Ishizaki et al., 2005; Ishizaki et al., 2006; Peng et al., 2015). The induction of these pathways is regulated by SnRK1 signaling, which activates a broader energy conservation program in response to carbon limitation (Dietrich et al., 2011; Pedrotti et al., 2018). These findings underscore the physiological importance of maintaining a robust, thermo-stable core of mitochondrial enzymes to support respiratory metabolism during abiotic stress.

In contrast, Pentatricopeptide Repeat (PPR) proteins involved in mitochondrial RNA editing consistently displayed low melting temperatures and were largely absent from the high-stability fractions. PPR proteins bind specific mitochondrial transcripts, and their stability *in vitro* may depend on the presence of substrate RNA. We therefore hypothesize that, *in vivo*, PPR proteins are stabilized upon mRNA binding and may be targeted for degradation when editing demand is low. Under hyperthermic or severe stress conditions that exceed the stability of unbound PPR proteins, a decline in RNA editing could lead to the accumulation of unedited respiratory complex subunits. Such defective subunits may fail to assemble properly, triggering reactive oxygen species (ROS) production and amplifying stress-responsive signaling cascades (Waltz et al., 2020).

Our findings are well in line with the results of a recent study on the thermal tolerance of the Arabidopsis proteome using CESTA (cellular thermal shift assay; Lyu et al., 2023)). While for our approach we isolated mitochondrial proteins to focus on interactions with cysteine as a ligand, CESTA works with intact cells preserving the diverse protein–protein and protein-metabolite interactions in the native cellular context. Both studies detected high thermal stability in mitochondrial dehydrogenases involved in the TCA cycle and amino acid metabolism. The extraordinary thermal vulnerability of mitochondrial RNA editing factors we observed in the protein extract also became apparent in the cellular context. Several PPR proteins were included in the 10% of the proteins with the lowest Tm within the CESTA study. In general, the proteostasis network, including aminoacyl-tRNA biosynthesis, protein translation initiation, and protein folding, was among the most vulnerable targets of heat stress in cells, and a suppression of protein synthesis by heat stress could be experimentally verified (Lyu et al., 2023). Overall, the comparison underscores that TPP on isolated mitochondrial proteins can provide valuable insights into intrinsic protein stability, even though *in vivo*, individual enzymes might be stabilized due to metabolon formation and protein-metabolite interactions that enhance thermal resilience.

### Thermal proteome profiling identifies mitochondrial cysteine aminotransferases

This study demonstrates that thermal proteome profiling, which was originally developed for drug target and off-target discovery in biomedical research (Savitski et al., 2014), is a powerful tool to identify plant enzymes metabolizing a compound of interest. By monitoring shifts in thermal stability upon L-cysteine treatment, we successfully re-identified three known mitochondrial cysteine-metabolizing enzymes, validating our approach as an effective screening tool. Encouraged by these positive controls, we targeted the long-hypothesized aminotransferase catalyzing the initial transamination of cysteine to 3-mercaptopyruvate (Höfler et al., 2016). Indeed, we could experimentally validate the interaction of cysteine with three mitochondrial aminotransferases showing a melting point shift in TPP. Activity tests confirmed that Asp-AT and Ala-AT can use cysteine as a substrate and catalyze the first step of the mitochondrial cysteine catabolic pathway. These results suggest that a dedicated cysteine aminotransferase is not required, but endogenous promiscuous aminotransferases can accommodate cysteine. This finding aligns with observations in mammalian mitochondria, where aspartate aminotransferase has similarly been implicated in cysteine transamination (Miyamoto et al., 2014). To our knowledge, no enzyme exclusively dedicated to cysteine transamination has been described in eukaryotes to date.

Beyond identifying candidate enzymes, the melting curves provided by TPP could potentially reveal some additional information on the mode of the enzyme-metabolite interactions. Aminotransferases require pyridoxal phosphate (PLP) as a cofactor. In the resting state, PLP is covalently linked to the enzyme via a Schiff base linkage formed by the condensation of its aldehyde group with the ε-amino group of a lysine residue at the active site (Kirsch et al., 1984). Upon substrate binding this internal aldimine is replaced by an external aldimine bond resulting in pyridoxamine-5’-phosphate (PMP), which is no longer covalently linked to the enzyme. In the absence of the second substrate the cofactor will eventually defuse off compromising the enzyme structure. The pronounced destabilization of Asp-AT, GABA-AT, and alanine-glyoxylate aminotransferase in the presence of cysteine likely reflects this cofactor release upon formation of a stalled external aldimine. In the case of GABA-AT we did not detect any cysteine aminotransferase activity, so the mechanism of inhibition might involve binding of cysteine at the active site without catalysis. Our data indicate a competitive character of the inhibition which would be in line with this mode of action. Conversely, Ala-AT’s stabilization by cysteine hints at an unusually strong non-covalent affinity for its cofactor–substrate complex, meriting further structural and spectroscopic investigation (Fig. 4).

### Metabolic integration of mitochondrial cysteine aminotransferases

Our study identified two mitochondrial aminotransferases able to use cysteine as an alternative substrate. Kinetic parameters place Ala-AT in a physiologically relevant role. Its K_m_ for cysteine (0.77 mM) is three-fold lower than for alanine and close to the physiological range of cysteine concentrations reported under normal and stress conditions (Heeg et al., 2008; Moormann et al., 2025; Watanabe et al., 2008). Moreover, both mitochondrial Ala-AT isoforms are upregulated by cysteine feeding in Arabidopsis seedlings, supporting their *in vivo* function in cysteine catabolism (Moormann et al., 2025). For Asp-AT we determined a comparatively high K_m_ of 15 mM with cysteine indicating that *in vivo* its activity will strongly depend on the cysteine concentration in the mitochondrial matrix.

Although aminotransferase activities with cysteine were lower than with canonical substrates, this may reflect metabolic flux distributions: alanine and aspartate are directly linked to central carbon metabolism via transamination to pyruvate and oxaloacetate, respectively. In contrast, cysteine catabolism via 3-mercaptopyruvate might be relevant for protein persulfidation (Pedre et al., 2023) and for the removal of accumulating cysteine during stress, while otherwise alternative branches in mitochondrial cysteine metabolism such as FeS synthesis or cyanide detoxification (Fig. 1) are most likely quantitatively more relevant. There could also be some presently unknown regulatory mechanism shifting the substrate preferences and catalytic efficiencies of the aminotransferases *in vivo*.

Co-expression profiles provide some first hints on the metabolic and physiological context of the different steps and variants of the mitochondrial cysteine catabolic pathway. Asp-AT and STR1 share an expression profile with core mitochondrial pathways, the TCA cycle and the respiratory chain, and are thus universally expressed. It is tempting to speculate that these two enzymes might also participate in functions of general importance, which could be related to the production of persulfides for signaling. In contrast, ETHE1 and GABA-AT are clearly stress responsive. Their expression profiles overlap with several enzymes from lysine and branched-chain amino acid catabolism that are also localized in the mitochondria and typically strongly induced by energy deficiency stress via SnRK1 signaling (Heinemann & Hildebrandt, 2021). In these stress conditions massive protein degradation releases high amounts of all proteinogenic amino acids so that the intracellular concentrations of those amino acids that are normally low abundant massively accumulate in the free pool (Heinemann et al., 2021). Cysteine belongs to this group of low abundant amino acids together with lysine and the branched-chain amino acids indicating that their catabolism might be regulated in a concerted fashion. The mitochondrial cysteine catabolic pathway we reconstruct here represents the only known route in plants that oxidizes the thiol moiety of cysteine instead of releasing hydrogen sulfide, a potent and potentially damaging gasotransmitter (Cooper & Brown, 2008). Thus, a physiological role in mitigating toxic amino acid accumulation during stress seems reasonable. The non-protein amino acid GABA also rapidly accumulates in plant tissues in response to biotic and abiotic stress. It enhances stress tolerance by regulating osmotic balance, improving antioxidant defense, and serving as a signaling molecule (Clark et al., 2009; Kinnersley & Turano, 2000). Our findings demonstrate that GABA-AT binds cysteine, most likely at the active site, without catalyzing its transamination. This is in line with a previous report that had tested cysteine and several other amino acids as substrates for GABA-AT (Clark et al., 2009). Instead, cysteine acts as an inhibitor of GABA-AT and thus its accumulation might serve as a regulatory brake on GABA turnover during acute stress. A high abundance of GABA-AT allows rapid removal of GABA after stress release providing significant amounts of anaplerotic carbon to the TCA cycle via the GABA shunt (Kinnersley & Turano, 2000).

## Conclusions

This study establishes thermal proteome profiling (TPP) as a valuable tool for identifying plant enzymes metabolizing a specific substrate of interest. By applying TPP to Arabidopsis mitochondrial proteins, we identified aminotransferases capable of initiating cysteine catabolism through transamination to 3-mercaptopyruvate. These findings enabled the reconstitution of a complete mitochondrial cysteine degradation pathway *in vitro*. The enzymatic properties and expression patterns of alanine aminotransferase and aspartate aminotransferase suggest distinct regulatory and physiological roles under stress conditions. Furthermore, cysteine’s inhibitory effect on GABA aminotransferase points to a potential regulatory interaction during metabolic adaptation. Beyond that, the TPP dataset represents a valuable resource for identifying potential targets of cysteine-mediated metabolic regulation, including transporters, RNA editing factors, and respiratory chain components. This aspect is especially relevant in the context of stress signaling, as cysteine has been implicated as a metabolic signal during both abiotic and biotic interactions (Heinemann et al., 2021; Moormann et al., 2025; Romero et al., 2014). Given the prevalence of allosteric regulation in amino acid biosynthetic pathways, our findings highlight the potential for cysteine to modulate enzyme activity through direct binding. Together, these insights deepen our understanding of mitochondrial sulfur metabolism and its potential integration into plant stress responses and metabolic signaling.

## Materials & Methods

### Plant material and protein extraction

Mitochondria were isolated from an *Arabidopsis thaliana* Col-0 cell suspension culture according to the procedure published in (Werhahn et al., 2001). The mitochondrial pellets were resuspended in 1 ml homogenization buffer (50 mM Tris (pH 7.5), 10 mM KCl, 3 mM EGTA, 0.4% NP-40) per 0.1 g pellet. The mild detergent NP-40 was included to extract membrane-bound proteins (e.g. receptors, (Reinhard et al., 2015). The mitochondrial fractions were incubated on ice for 30 min, ruptured in a Potter homogenizer, frozen at -20°C and stored until the thermal treatment experiment.

### Temperature gradient precipitation

We used approx. 10 ml of mitochondrial protein extract (∼1 µg µl^-1^) to perform a thermal proteome profiling experiment. With this amount we could cover three replicates for ten different temperatures of a control and an L-cysteine treated variant. In detail, the mitochondrial extract was split in half and was either incubated with 5 mM L-cysteine or water for 10 min on a shaker at room temperature. The treated extracts were then aliquoted (150 µl) into PCR-tubes for the temperature treatment in a thermocycler. Three aliquots per treatment and temperature were prepared and incubated for 10 min at either 25°C, 30°C, 35.8°C, 40.7°C, 44°C, 50°C, 55.2°C, 61.4°C, 66°C or 70°C. The treated and heated extracts were then transferred into 1.5 mL reaction tubes and centrifuged at 20,000 x g for 20 min to pelletize the potentially denatured proteins. Supernatants were collected and protein concentrations were quantified via Bradford assay (Thermo Fisher Scientific). Subsequently, 1 µg bovine serum albumin (BSA) was added to each sample as internal standard to preserve the information of different protein concentrations. Eventually, the protein samples were concentrated in a vacuum concentrator for further processing.

### Protein digestion and sample preparation for proteome analysis via mass spectrometry

We used a modified version of the “solvent precipitation, single-pot, solid-phase-enhanced sample preparation (SP4) protocol of (Johnston et al., 2022). The concentrations of the protein samples were adjusted to 1 µg/µL by adding individual volumes of SDT buffer (4% SDS, 0.1 M dithiothreitol, 0.1 M Tris pH 7.6, (Mikulášek et al., 2021). The TPP extracts were incubated at 60°C for 30 min to solubilize, denature and reduce the proteins. Samples were then sonicated for 10 min and centrifuged at 20,000 ×g for 10 min. Thirty microliters of the supernatant were transferred to new tubes, mixed with 7.5 µL iodoacetamide (IAM, 0.1 M) and incubated for 30 min in the dark in order to alkylate the reduced disulfide bridges. Then 2 µL DTT (0.1 M) was added to neutralize excess amounts of IAM.

Preparation of the glass beads/ACN suspension, protein precipitation and washing steps were performed as described in (Johnston et al., 2022). We used approx. 400 µg of glass beads per sample (approx. 30 µg protein). The purified proteins were digested with 0.5 µg trypsin (mass spectrometry grade, Promega) for 16 h on a heated shaker at 37°C at 1000 rpm. The peptide-containing supernatants were collected in low peptide binding tubes. The glass beads were rinsed in 60 µL ammonium bicarbonate (50 mM) to recover remaining peptides. Eluates were combined and acidified with 1 µL formic acid (FA). The peptides were desalted on 50 mg Sep-Pak tC18 columns (WAT054960, Waters) and quantified using the Pierce Quantitative Colorimetric Peptide Assay Kit (Thermo Fisher Scientific). The samples were finally diluted to a final concentration of 400 ng μl^−1^ in 0.1% FA.

### Quantitative proteomics by shotgun mass spectrometry (LC-MS/MS)

400 ng of peptides were injected via a nanoElute2 UHPLC (Bruker Daltonic) and separated on an analytical reversed-phase C18 column (Aurora Ultimate 25 cm x 75 µm, 1.6 µm, 120 Å; IonOpticks). Using a multistage linear gradient (eluent A: MS-grade water containing 0.1% formic acid, eluent B: acetonitrile containing 0.1% formic acid, gradient: 0 min, 2% B; 54 min, 25% B; 60 min, 37% B; 62 min, 95% B; 70 min, 95% B), peptides were eluted and ionized by electrospray ionization using a CaptiveSpray 2 source with a flow rate of 300 nL/min. The timsTOF-HT mass spectrometer followed a data-dependent acquisition parallel accumulation– serial fragmentation (DDA-PASEF) method, covering an ion mobility window of 0.7–1.5 V s/cm^2^ with 4 PASEF ramps, targeting an intensity of 14,500 (threshold 1,200) with a cycle time of ∼0.53 s. Ion mobility spectrometry–MS/MS spectra were analyzed with MaxQuant (Cox and Mann, 2008) using default search parameters and the TAIR10 database for protein identification. Additionally, the calculation of label-free quantification (LFQ) values and intensity-based absolute quantification (iBAQ) values for the identified proteins were enabled.

### Thermal proteome profiling data analysis

For thermal proteome profiling data analysis, we developed a custom R script (MeCuP, available on demand) which enables automated processing and visualization based on input from MaxQuant. Input data is structured according to the ARC directory format and includes MaxQuant output files (proteingroups.txt with iBAQ values), sample annotation (.csv format), and optional protein annotations. The script normalizes protein abundance data based on BSA-spiked internal standards and applies a starting point (SP) mean normalization at the lowest temperature (e.g. 25°C or user-defined). Normalization can be performed against control, treatment-specific or both respectively SP values. Samples can be filtered based on BSA abundance and the number of valid data points at SP, ensuring only reliable melting curves are included in the analysis. Key user-defined parameters include thresholds for BSA detection, SP data filtering, and minimum valid datapoints. Proteins failing these criteria are excluded and listed in the output. Filter effects can be explored separately. Melting points (Tm50) are calculated from LOESS-smoothed curves as the temperature corresponding to 50% soluble fraction. Visual outputs include protein melting curves, with optional emphasis on non-overlapping curves between control and treatment. Final outputs include customizable .xlsx files with legends, plots, exclusion logs, and a result table with calculated Tm50 values per protein and condition.

### Cloning of Arabidopsis cysteine binding candidates

Full-length cDNA for candidate enzymes (STR1: AT1G79230; ETHE1: AT1G53580 Ala-AT: AT1G17290; Asp-AT: AT2G30970; GABA-AT: AT3G22200) were cloned using RT-PCR from total RNA isolated from mature Arabidopsis leaves. Total RNA was extracted using Monarch total RNA Miniprep kit (New England Biolabs) following the manufacturer’s protocol. The total RNA was also additionally treated with DNase1 (New England Biolabs) as instructed by the manufacturer. The isolated RNA was quantified using nanophotometer (NanoPhotometer® N60, Implen) and was stored in -80°C immediately. GoScript Reverse Transcriptase (Promega) was used to synthesise cDNA from 5 μg of RNA following manufacture’s protocol with the exception of using anchored oligo(dT)20 VN primers instead of oligo(dT)15 primers. Additionally, RNasin (Promega) was also added in the reaction mix. Synthesised cDNA was used for subsequent gene specific PCR’s using Phire Hot Start II DNA Polymerase (Thermo Fisher Scientific) with gene specific primers (Suppl. Table S1). The mitochondrial targeting sequences for all enzymes were removed at their cleavage sites and exchanged with Met codon. The primers contained restriction enzyme sites along with compatible overhangs for homology based directional cloning into pET28a(+) (Novogen) vector. The amplified sequences were then ligated into pET28a(+) (Novogen) plasmids using NEBuilder HF DNA assembly kit (New England Biolabs) following the manufacturers’ instructions. The ligated plasmids were transformed into NEB 5-alpha competent *Escherichia coli* cells (New England Biolabs) and grown in Luria-Bertani (Miller) (LB) (Sigma-Aldrich) media supplemented with 50 μg ml−1 kanamycin. Positive clones were screened and sequence verified through sanger sequencing (Microsynth Seqlab).

### Protein overexpression and purification

The verified plasmids were transformed into expression capable BL21(DE3) competent *E. coli* (New England Biolabs) cells. For some candidates (GABA-AT and ETHE1) the competent BL21(DE3) *E. coli* harboured an additional plasmid-pG-KJE8 (Takara) containing coding sequences for molecular chaperons for assisted folding. Overexpression of the recombinant *E. coli* (OD_600 nm_ ≈ 0.4) proteins was induced using 1 mM isopropyl β-D-1-thiogalactopyranoside (IPTG) (Thermo Fisher Scientific). For cells harbouring chaperons, addition of 10 ng ml^-1^ Tetracycline and 0.5 mg ml^-1^ of L-Arabinose preceded IPTG to allow for sufficient production of chaperons required for assisted folding. Additionally, cultures of candidate aminotransferases (ALA-AT, Asp-AT and GABA-AT) were supplemented with 0.1 mM of pyridoxal 5’-phosphate (PLP) to enhance folding and stability of protein. The cultures were allowed to grow at 18°C overnight for 16h with gentle agitation in baffled flasks.

The cells were harvested by chilling the cultures on ice for 15 min followed by centrifugation at 16,000 x g for 20 min at 4°C. The pellets were suspended in ice cold lysis buffer (50 mM sodium phosphate, 500 mM NaCl, 10 mM Imidazole, 250 mM Sucrose, 5% Glycerol, pH 8.0 containing 2 mg ml^-1^ of Lysozyme, 10 units of DNase1 and RNase1) and incubated at RT for 1 hour. Lysis buffer is also supplemented with EDTA-free protease inhibitor (Roche). GABA-AT was processed differently by using 50 mM Tris(hydroxymethyl)methyl-3-aminopropanesulfonic acid (TAPS) instead of sodium phosphate at pH 9, the rest of buffer components remained the same. The cells were sonicated for 2 minutes with a pulse cycle of 20 sec ON and 30 sec OFF at 70% amplitude. After sonication the supernatant was harvested by centrifugation at 16,000 x g at 4°C for 15 min. The supernatant was passed through 0.45 μm filter to remove any cell debris.

Purification of the recombinant enzymes were carried out using affinity chromatography on 5 ml nickel-nitrilotriacetic acid-agarose (NiNTA) columns (Cytiva) coupled to an ÄKTA go system (Cytiva). 50 ml of clarified supernatant was injected into a pre-equilibrated column at a flowrate of 1 ml min^-1^ followed by washing with binding buffer for 5 CV (50 mM sodium phosphate/ 50 mM TAPS, 500 mM NaCl, 250 mM sucrose, 5% glycerol, pH 8.0/ 9.0). The bound proteins were eluted using a gradient of imidazole (0-500 mM) for 10 CV at a flowrate of 1 ml min^-1^ while collecting 2 ml fractions. The purified protein fractions were pooled and concentrated to 500 μl using Amicon 10K Ultra Centrifugal Filters (Merck). Concentrated samples were subjected to size exclusion chromatography (SEC) using Superdex 200 increase 10/300 (Cytiva) column coupled to an ÄKTA go system (Cytiva). The proteins were separated based on their molecular weights using 2 CV of SEC running buffer (50 mM sodium phosphate, 300 mM NaCl, 10% glycerol, pH 8.0 for ETHE1, STR1, Ala-AT and Asp-AT; 50 mM TAPS, 300 mM NaCl, 10% glycerol, pH 9.0 for GABA-AT) at a flowrate of 0.1 ml min^-1^. 500 μl fractions were collected and fractions containing distinct peaks were screened through 10% (Ala-AT, Asp-AT, GABA-AT) and 15% (STR1 and ETHE1) polyacrylamide gels and visualised using fast Coomassie stain (Serva). The same were also verified using western blot using anti-His mouse monoclonal antibody (Cell Signalling Technologies) and anti-mouse IgG HRP-linked antibody (Cell Signalling Technologies). The purified enzymes fractions corresponding to the right electrophoretic mobility were stored at -20°C in 50% glycerol (v/v). These fractions were quantified using Bradford assay (Thermo Fisher Scientific) and used for enzymatic tests.

### Enzyme activity tests

Activity assays for Ala-AT and Asp-AT contained 50 mM 4-(2-hydroxyethyl)-1-piperazineethanesulfonic acid (HEPES) pH 8.0, 50 µM pyridoxal-5-phosphate (PLP) and 0.25 – 0.3 µg ml^-1^ protein for the standard reaction and 200 – 250 µg ml^-1^ protein for reactions with cysteine as amino donor. If not indicated otherwise, 9 mM alanine and 12 mM aspartate were used as amino donors for Ala-AT and Asp-AT reactions, respectively, while 6 mM 2-oxoglutarate was used as amino acceptor. K_m_ and V_max_ of the respective substrates were determined using 0.25 – 12 mM alanine and 0.125 – 1.5 mM cysteine for Ala-AT; 0.5 – 21 mM aspartate and 3 – 80 mM cysteine for Asp-AT and 0.1 – 6 mM 2-oxoglutarate for both Ala-AT and Asp-AT. Standard activity assays for GABA-AT contained 50 mM tris(hydroxymethyl)methyl-3-aminopropanesulfonic acid (TAPS) pH 9.0, 50 µM PLP and 3.7 - 5.9 µg ml^-1^ protein. Inhibitory effects of cysteine were determined using 0, 1, 2 and 5 mM cysteine. Forward reactions contained 1 – 8 mM GABA and 6 mM pyruvate. Reverse reactions contained 1 – 7.5 mM alanine and 0.5 mM succinic semialdehyde. Reactions that generated pyruvate or oxaloacetate were coupled to NADH-dependent lactate dehydrogenase (LDH, 0.3 U or 5 µg ml^-1^, pig heart, Roth) or malate dehydrogenase (MDH, 2.4 U or 10 µg ml^-1^, pig heart, Roche), respectively. Continuous assays contained 0.3 mM NADH and the reactions were initiated by the addition of an appropriate amino donor or amino acceptor. The rate of each reaction was monitored as the change in NADH concentration at 340 nm using the spectrophotometer Multiskan Skyhigh (Thermo Fisher Scientific) equipped with temperature control. In assays that were not coupled to LDH or MDH, the formation of amino acids was monitored using reverse-phase high performance liquid chromatography (HPLC). The discontinuous assays were stopped by transferring aliquots into 0.1 M HCl. Samples were neutralized by the addition of 0.5 M potassium borate buffer pH 11. Amino acids were quantified using the Agilent 1260 Infinity II HPLC System (Agilent) and pre-column derivatization with o-phthaldialdehyde (OPA) and fluorenylmethoxycarbonyl (FMOC) based on the application note “Automated amino acid analysis using an Agilent Poroshell HPH-C18 Column” by Agilent. Peaks were evaluated and quantified using OpenLabCDS software (Agilent). Reaction rates were determined based on at least 3 time points. All assays were conducted in 30 °C and in triplicates. Kinetic parameters and inhibitory constants were calculated using the non-linear regression least squares fit analysis “Michealis-Menten” and “Competitive inhibition”, respectively, in GraphPad Prism (Version 10.4.2 for Windows, GraphPad Software).

### Pathways Reconstruction

All reactions contained 50 mM HEPES pH 8.0, 1 mM glutathione and 1.1 µg ml^-1^ ETHE1. Assays for ETHE1 reactions were started by adding 10 µl of a saturated sulfur solution (in acetone). Assays for combined STR1 and ETHE1 reactions contained additional 7.1 µg ml^-1^ STR1 and were started by adding 1 mM 3-mercaptopyruvate. Assays for full pathway reconstitution contained additional 7.1 µg ml^-1^ STR1, 50 µM PLP, 1 mM 2-oxoglutarate, 63 µg ml^-1^ Asp-AT or 99 µg ml^-1^ Ala-AT and were started by adding 1 mM cysteine. Oxygen content was continuously measured using a clark type oxygen electrode (model DW1, Hansatech Instruments Ltd). All assays were performed in triplicates at 30 °C and reaction rates were determined from the linear phase of oxygen depletion over 20 s. After 50 nmol of oxygen was consumed, samples for thiol quantification were transferred into derivatization buffer (1.5 mM bromobimane; 32% (v/v) acetonitrile; 10.3 mM EDTA and 10.3 mM HEPES pH 8) and incubated for 10 min on ice before the addition of methanesulfonic acid (final conc. 15.9 mM). Samples were diluted and measured using an Agilent 1260 Infinity II HPLC (Agilent) by fluorescence detection (ex. 380 nm; em. 480 nm). Peaks were evaluated and quantified using OpenLabCDS software (Agilent). Assays for oxygen content traces (Fig. 6 B, C, D, E) were started by the addition of 1.1 µg ml^-1^ ETHE1 and were repeated at least 3 times with similar results.

## Data availability

The mass spectrometry proteomics data have been deposited to the ProteomeXchange Consortium (http://proteomecentral.proteomexchange.org) via the PRIDE partner repository (Perez-Riverol et al., 2022) with the dataset identifier PXD064096.

## Acknowledgments

We are grateful to Hans-Peter Braun for valuable discussions on TPP, for hosting part of the group in his lab, and for supporting us with mitochondrial preparations. We thank Christina Mack and Dagmar Lewejohann for skillful technical assistance.

