## Supplementary Figures S1-6 for "Thermal proteome profiling identifies mitochondrial aminotransferases involved in cysteine catabolism via persulfides in plants"

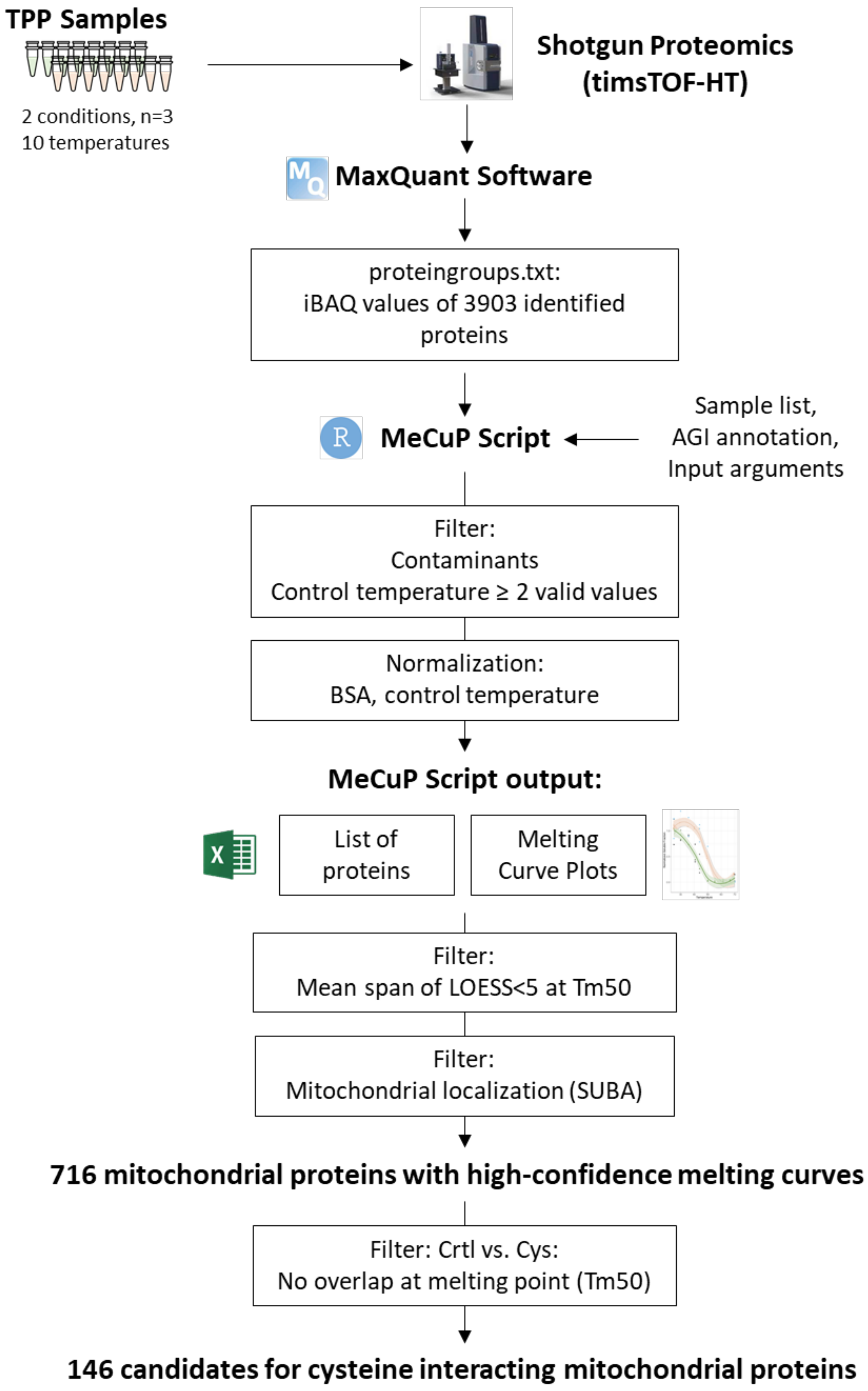


**Suppl. Figure S1:** Thermal Proteome Profiling (TPP) data processing workflow

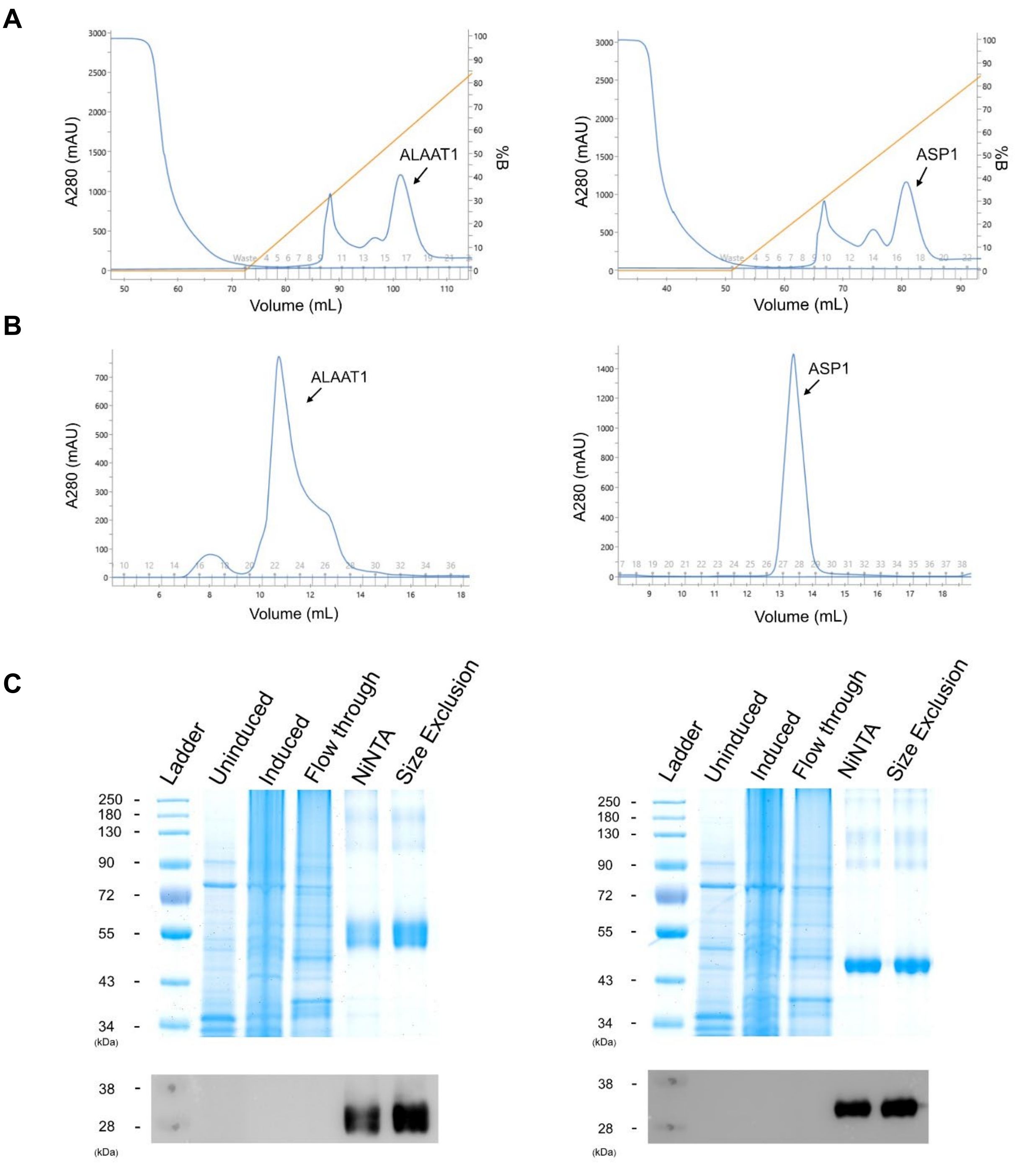


**Suppl. Figure S2:** Purification of recombinant AlaAT and AspAT

**A** UV chromatograms showing purification peaks for different eluting proteins during gradient elution from NiNTA column. ALAAT and ASP. **B** UV chromatograms showing singular peaks of size excluded fractions for ALAAT and ASP. **C** SDS-PAGE gel (Lanes 1-6) showcasing, Marker (in kDa), Uninduced, Induced, Flowthrough along with NiNTA purified and Size exclusion fractions. ALAAT and ASP correspond to their MW of 54.7 kDa and 45.8 kDa. A Corresponding western blot confirms the purification and presence of our recombinant protein purified from *E. coli* cultures.

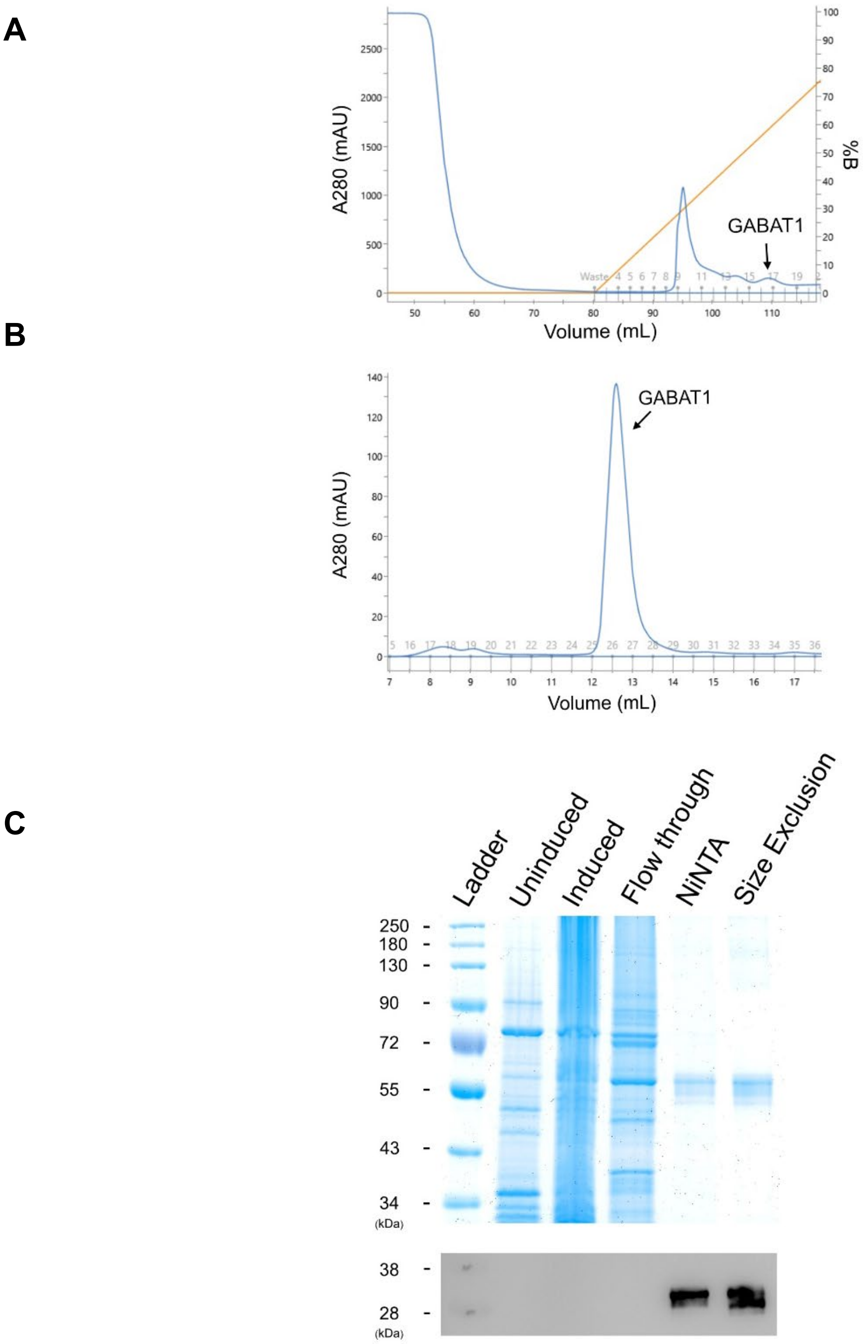


**Suppl. Figure S3:** Purification of recombinant GABA-AT

**A** UV chromatograms showing purification peaks for different eluting proteins during gradient elution from NiNTA column. GABAT. **B** UV chromatogram showing singular peaks of size excluded fractions for GABAT. **C** SDS-PAGE gel (Lanes 1-6) showcasing, Marker (in kDa), Uninduced, Induced, Flowthrough along with NiNTA purified and size exclusion fractions. GABAT correspond to MW of 53.3 kDa. A Corresponding western blot confirms the purification and presence of our recombinant protein purified from e coli cultures.

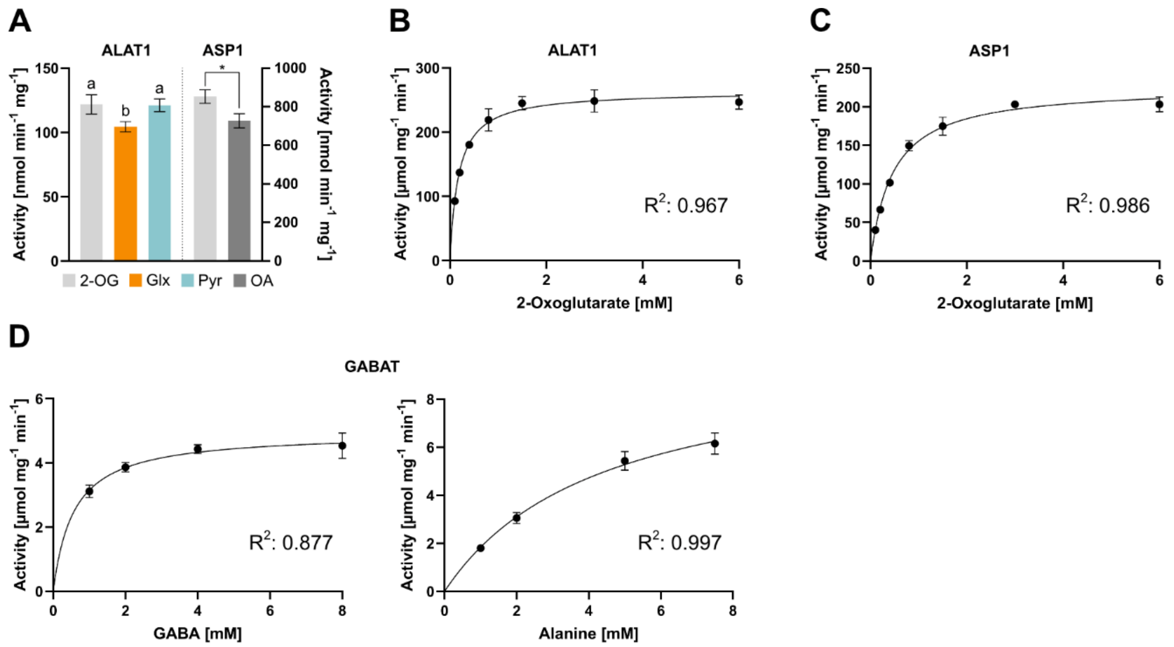


### Fig. 5: Biochemical characterization of mitochondrial cysteine aminotransferase candidates

**A** Aminotransferase activities of Ala-AT with Ala as amino donor and AspAT with Asp as amino donor in combination with different ketoacid substrates **B** Michaelis Menten kinetics for alanine aminotransferase 1 (AT1G17290) with L-Ala as fixed and 2-oxoglutarate as variable substrate. **C** Michaelis Menten kinetics for aspartate aminotransferase 1 (AT2G30970) with L-Asp as fixed and 2-oxoglutarate as variable substrate. **D** Michaelis Menten kinetics for GABA aminotransferase 1 (AT3G22200) with GABA and Ala as variable substrate.

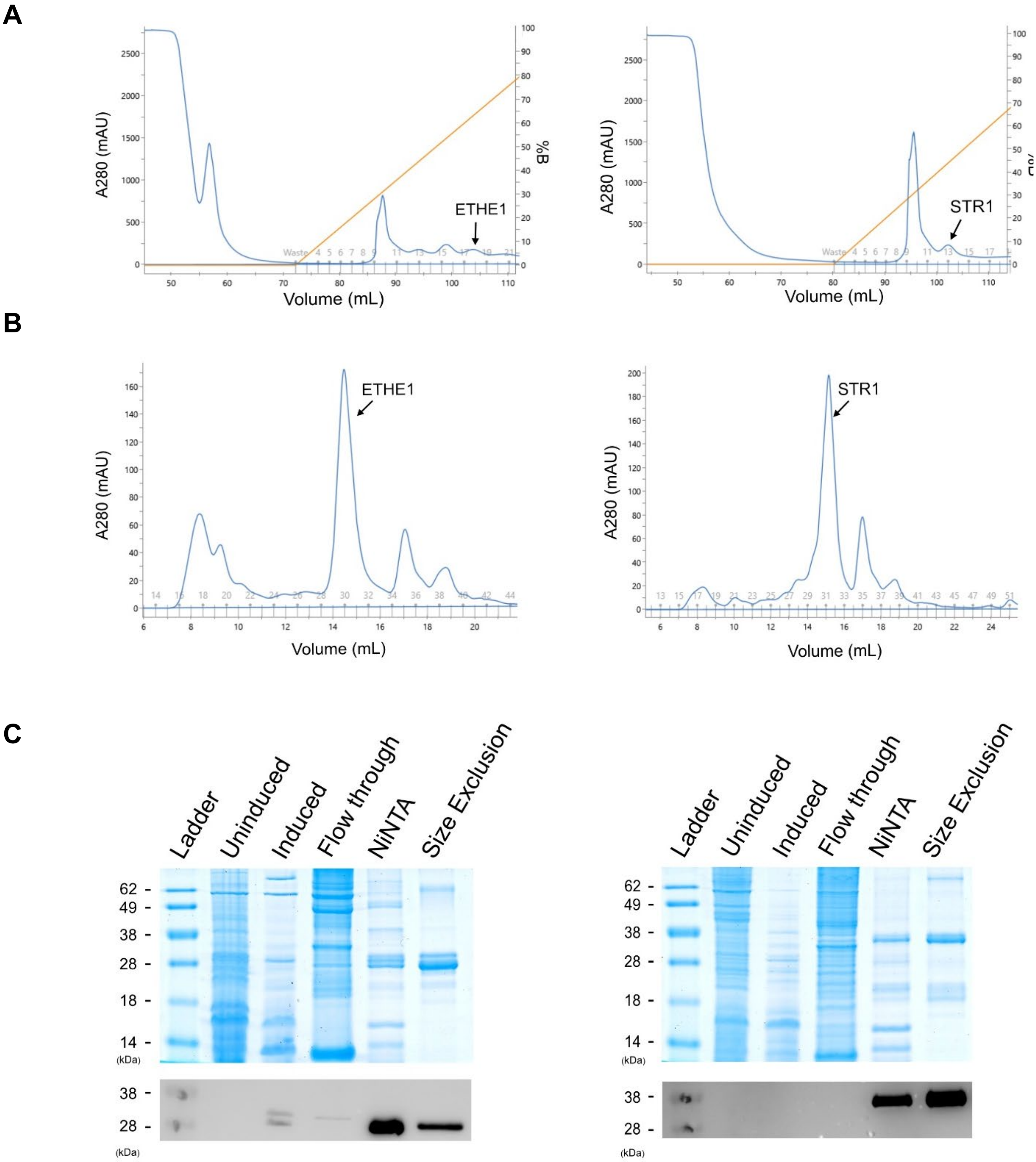


**Suppl. Figure S5:** Purification of recombinant ETHE1 and STR1

**A** UV chromatograms showing purification peaks for different eluting proteins during gradient elution from NiNTA column. ETHE1 and STR1. **B** UV chromatogram showing singular peaks of size excluded fractions for ETHE1 and STR1. **C** SDS-PAGE gel (Lanes 1-6) showcasing, Marker (in kDa), Uninduced, Induced, Flowthrough along with NiNTA purified and Size exclusion fractions. ETHE1 and STR1 correspond to their MW of 28.9 kDa and 36.6 kDa. A Corresponding western blot confirms the purification and presence of our recombinant protein purified from *E. coli* cultures.

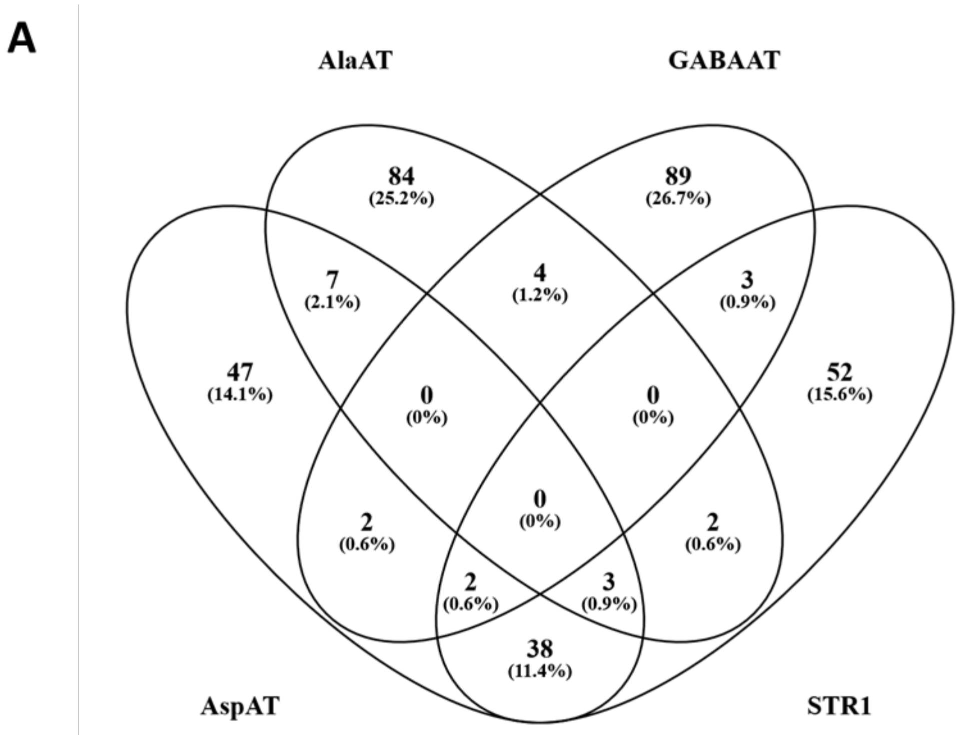


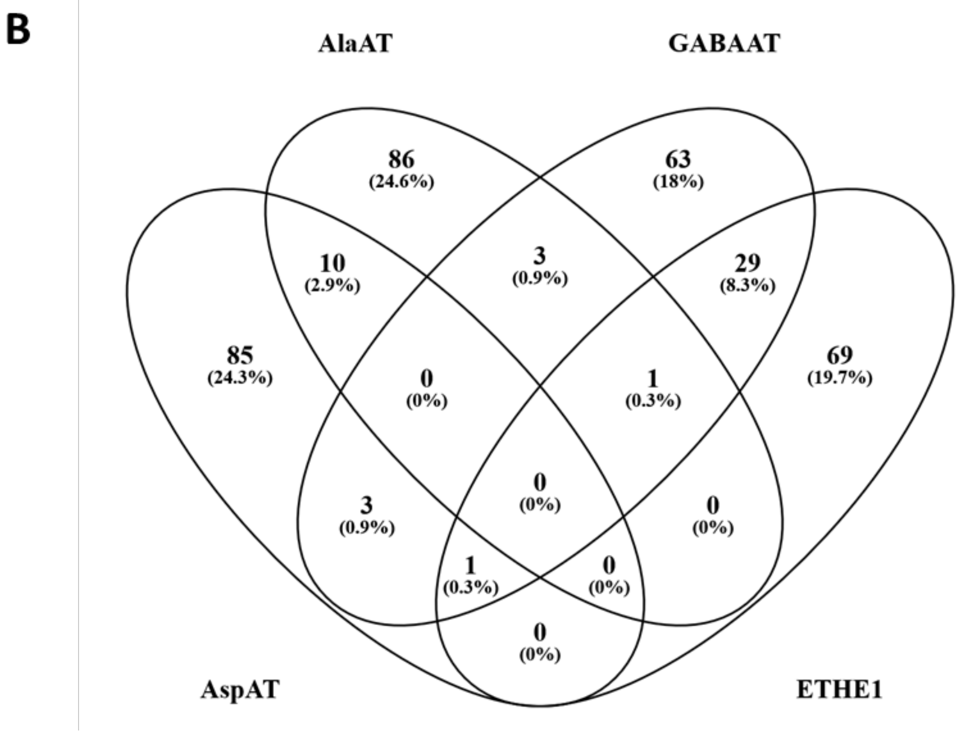


### Suppl. Figure S6: Overlap between coexpression profiles of the enzymes involved in mitochondrial cysteine catabolism

To get a first impression about the metabolic integration of the three aminotransferases in mitochondrial sulfur metabolism as well as the physiological context of these networks we compared co-expression profiles available on ATTED (https://atted.jp/). The top 100 coexpressed genes for the individual enzymes are listed in Suppl. Dataset S2.
